# Unique K-mer sequences for validating cancer-related substitution, insertion and deletion mutations

**DOI:** 10.1101/2020.06.20.163113

**Authors:** HoJoon Lee, Ahmed Shuaibi, John M. Bell, Dmitri S. Pavlichin, Hanlee P. Ji

**Affiliations:** Division of Oncology, Department of Medicine, Stanford University School of Medicine, Stanford, CA, 94305, United States; Stanford Genome Technology Center, Stanford University, Palo Alto, CA, 94304, United States

## Abstract

The cancer genome sequencing has led to important discoveries such as identifying cancer gene. However, challenges remain in the analysis of cancer genome sequencing. One significant issue is that mutations identified by multiple variant callers are frequently discordant even when using the same genome sequencing data. For insertion and deletion mutations, oftentimes there is no agreement among different callers. Identifying somatic mutations involves read mapping and variant calling, a complicated process that uses many parameters and model tuning. To validate the identification of true mutations, we developed a method using k-mer sequences. First, we characterized the landscape of unique versus non-unique k-mers in the human genome. Second, we developed a software package, KmerVC, to validate the given somatic mutations from sequencing data. Our program validates the occurrence of a mutation based on statistically significant difference in frequency of k-mers with and without a mutation from matched normal and tumor sequences. Third, we tested our method on both simulated and cancer genome sequencing data. Counting k-mer involving mutations effectively validated true positive mutations including insertions and deletions across different individual samples in a reproducible manner. Thus, we demonstrated a straightforward approach for rapidly validating mutations from cancer genome sequencing data.

## INTRODUCTION

Next generation sequencing **(NGS)** analysis has been widely adopted in cancer research for identifying mutations and other genetic aberrations (1). For example, the Cancer Genome Atlas Project **(TCGA)** has relied on exome sequencing to identify numerous driver mutations in over 30 cancers. These catalogues of cancer genetic alterations provide insight to the underlying mechanisms of cancer and are used clinically for predicting response to certain therapies and have prognostic implications (2–4). However, the analysis of cancer genome sequencing data relies on human genome assemblies for reference alignment. Since mapping sequence reads enables the identification of mutations, their positions, and their allelic fractions, accurate variant analysis depends on alignment reference mapping accuracy.

Although cancer genome sequencing has become routine for biomedical research studies and diagnostic genetic testing of tumors, there are significant challenges in accurately identifying cancer mutations. The complexity of this task is evident in the discordant results produced by different mutation callers (5). The relatively sparse overlap among various mutations callers is a major dilemma and directly stems from the use of the human reference genome for sequence alignment (6). The human reference build is a static representation of sequences derived from the genomes of 13 individuals, encompasses only a small proportion of human genome diversity and lacks feature indicating structural complexity. The broad spectrum of novel and complex somatic alterations present in cancer genomes are often missed or misclassified due to these limitations of the reference genome. For example, short sequence reads unique to a specific tumor genome and containing a novel mutation may prove to be unmappable and thus are excluded from analysis. Unknown homologous or paralogous genes are similarly problematic due to uncertain mapping locations of the genomic reads.

A major challenge in alignment-based variant calling is the identification of insertions and deletions **(indels)**; this challenge is intrinsically related to alignment scoring metrics. Single nucleotide variants **(SNVs)**such as substitutions have a lower alignment penalty score than indels. As a result, reads with substitutions have higher alignment scores compared to reads with indels. Lowering the penalty for insertions and deletion does not resolve this issue (7). For example, we observed that modifying the penalty threshold resulted in the interpretation of true substitutions as 1-base indels whenever adjacent bases matched the alternate allele. Altering these settings leads to the calling of spurious indels for approximately 25% of the substitutions. Furthermore, variant calling programs such as GATK (8), MuTect2 (9) and VarScan2 (10) require statistical models to identify mutation events from mapped reads. Despite these programs’ ability to identify true positive mutations, there are issues with reproducibility when making comparisons among different callers or even when using the same caller repeatedly (11,12). For example, repeating the analysis for discovering mutations can lead to a different set of variant calls despite starting from the same data set of mapped reads.

For this study, we examined the properties of k-mers, short segments of DNA sequence, that include somatic mutations identified in cancer genome sequencing data. Our goal was to determine the k-mer properties of somatic mutations and to assess the properties of unique k-mers for validating true versus artifactual mutation calls. Previous studies have utilized k-mers to analyze sequences from organisms lacking a complete reference genome (13). K-mers are used in sequencing alignment programs such as BLAST, BLAT, BWA (14–16) and RNA-seq analysis (17,18). Some studies have used k-mers to genotype known variants (19) or capture reads for efficient target mutation validation from targeted sequencing (20). However, none of these previously published studies have conducted a thorough, systematic evaluation of the property of k-mers related to cancer mutations found in exome or whole genome sequence data.

For this analysis, we developed a computational tool, KmerVC, that determines the properties of k-mers such as their uniqueness in the human genome reference. Moreover, we determined which specific k-mer properties defined mutations as somatic. After obtaining k-mer counts, we used a binomial statistical test to validate cancer mutations and given mutation calls. Our results suggest that using k-mers without a conventional static reference has the potential for first pass mutation calling.

## MATERIAL AND METHODS

We implemented KmerVC as a Python 2.7 software package consisting of a command-line program, kmervc.py, and a reusable library, kmervclib. The github repository is open access at https://github.com/compbio/kmerVC. The KmerVC program requires three inputs: (i) a list of mutation calls of interest in VCF or BED file format (ii) the reference genome sequence, and (iii) the primary sequencing data in FASTQ or FASTA format.

### The overall analysis pipeline of KmerVC

The overall structure of KmerVC is outlined in **Figure 1**. Our tool has five steps; i) pre-processing to assess the uniqueness of a k-mer in the human genome, ii) counting k-mers in sequencing data, iii) retrieving expected k-mers from regions surrounding mutation calls of interest, iv) compiling k-mer counts related to called mutations from a variant caller, and v) validating true positive mutations.

**Figure 1.**
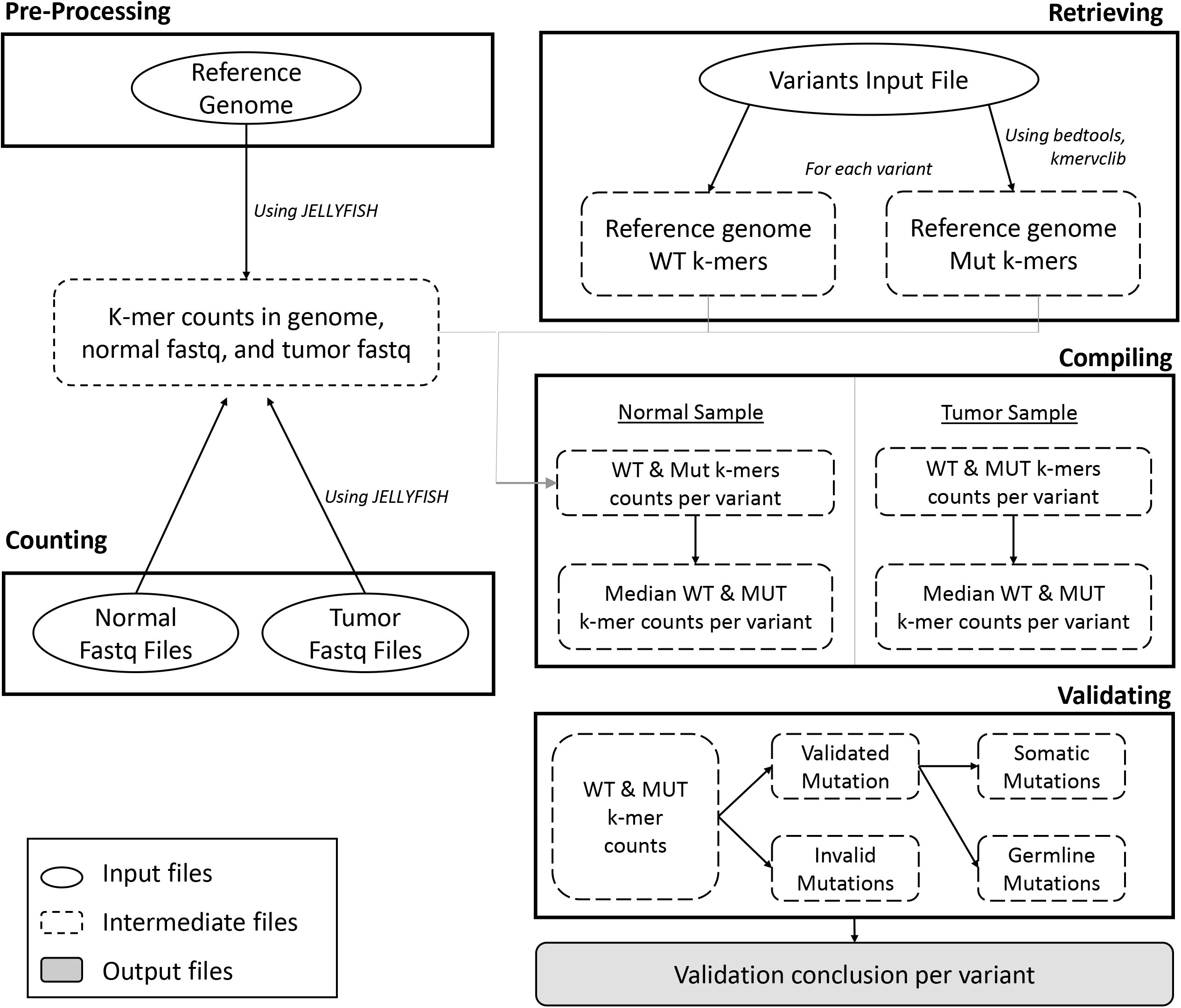
Overview of pipeline. (**a**) Preprocessing. (i) Determining the frequency of every distinct k-mer in the reference genome to ascertain its uniqueness. (ii) Determining the frequency of k-mers in normal and tumor FASTQ input files using JELLYFISH: a fast k-mer counting software. **(b)** Extraction. For all variants, we obtained the surrounding sequence region and generated a set of respective k-mers that include the target. Overlapping regions have the variants considered separately and consecutively accounted for in decomposition. Finally, non-unique normal and nonzero mutant k-mers are filtered from the sets. **(c)** Compilation. For all variants, we obtained the frequency of the corresponding k-mer set from the pre-processed counts dictionaries. **(d)** Validation. We assess if the variants are germline, somatic, or otherwise using a binomial test. For the binomial test, we utilize a sequencing error rate of 0.01 and an alpha of 0.01. We determine if the median count of wild-type and mutant k-mers are nonequivalent in the normal sample in the first test and if the median count of wild-type and mutant k-mers are nonequivalent in the tumor sample in the latter.

#### Pre-processing

With the default settings, we used JELLYFISH, a k-mer frequency counting software to obtain k-mer counts (21) using the command line call:

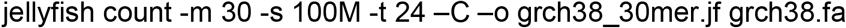

#### Counting k-mers

Accounting for multiple FASTQ input files, we obtain k-mer counts using the command line calls:

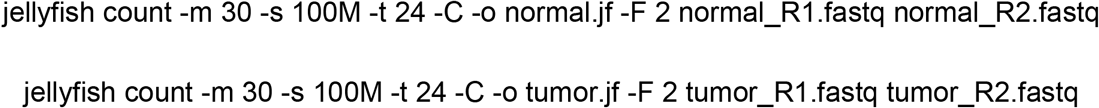

#### Retrieving expected k-mers

We extracted the sequence regions and generated a set of k-mers encompassing each mutation. This process occurs with a list of list of mutation coordinates used to generate a BED file. For a given mutation coordinate and a specific k-mer length denoted by *k*, each segment is defined by the mutation coordinate minus (*k – 1*) and plus *k*. With this BED file, we extract the corresponding sequences from the reference genome using the genomic analysis toolkit, BedTools. Then, we generated a FASTA file of the surrounding sequence regions with the command:

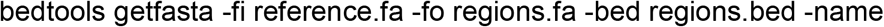

The BedTools getfasta functions through indexing the reference genome FASTA file and quickly identifying the specific sequence segment. We selected the use of Bedtools over another genome analysis toolkit Samtools, due to its faster runtime efficiency. Using the segment of sequence surrounding each variant, we determined whether the k-mers were wild-type, representing the reference versus being one derived from mutation. Wildtype k-mers were obtained by generating all k-mers that cover a segment containing the variant position of interest. Mutation-containing k-mers were generated: substituting the specified mutation at the given variant position and similarly generating all k-mers that include the mutation such as a substitution. Subsequently, we constructed a list of wild-type and mutant k-mers particular to each variant provided as input. This yielded a set of k-mers for each variant that we utilize to assess its validation.

Further, where two somatic mutations exist in one k-mer region, we examined two potential scenarios that would result due to the diploid chromosome: i) both mutations exist on the same chromosome, or ii) each of the mutations exist on different chromosomes. The mutation k-mers differ in these two cases and they are validated separately. In addition, we considered a scenario in which there are multiple mutations in a k-mer region. In this case, we deal with each pair of consecutive mutations as we would the case of two mutations in a region as just described. Cases in which more than two mutations are contained in a single k-mer region were discarded from further analysis and reported as such.

#### Compiling

We evaluated the total counts of unique k-mers in the reference genome per variant. To identify a variant given its surrounding k-mers, it is important that a portion of the k-mers are unique. We only proceeded with the statistical analysis of variants that had five or more unique k-mers in its surrounding region. We calculated the median count for each variant’s generated set of wild-type and mutant k-mers. We used the median value since it was more robust to sequencing errors and yielded more reliable results.

#### Validating

We performed statistical analysis using a binomial or Poisson test provided by the scipy.stats python package. First, we tested if a candidate variant is germline by determining the difference between wildtype and mutation k-mer counts in the normal sample. Thereafter, we determined if the variant is somatic using the difference between counts of wildtype and mutation k-mer counts in the tumor sample. We assumed a sequencing error rate of 1% and utilized an alpha value of 0.01. For the insertion/deletion, we fixed sequencing error rate as 0.01 regardless of their length, which is more stringent threshold compared to adjusted lower sequencing error rate for longer indels. These hyperparameters were tuned using simulated data although users can choose their own value. In addition, we applied a Bonferroni correction to the alpha value based on the number of tested variants. We validated a variant when the tests i) failed to reject null hypothesis (i.e. that it is wildtype) in the normal sample and ii) rejected the null hypothesis in the tumor sample, thereby qualifying it as a somatic mutation. In the case that both null hypotheses were rejected, the variant was determined to be germline. In all other cases, the variant was considered an artifact. We required that the count of unique wild-type k-mers and the count of unique mutant k-mers, which should not present in the genome, was a threshold of five or above. This threshold value ensured that the identified variants could be mapped back to the genome with accuracy and robustness. Calls passing this criterium were determined to be true positive somatic variants and thus validated.

We provide a validity assessment for each input variant regarding its validity as described among one of several categories **(Figure 2)**. A variant marked as ‘insufficient’ has not met the proper conditions regarding the count of surrounding unique wild-type and unique mutant k-mers. As a result, assessment of this variant and its validation is unreliable. A variant marked *SNP_affected* possesses a count of wild-type k-mers near 0 and thus, the k-mer properties are insufficient to proceed with the validation.

**Figure 2.**
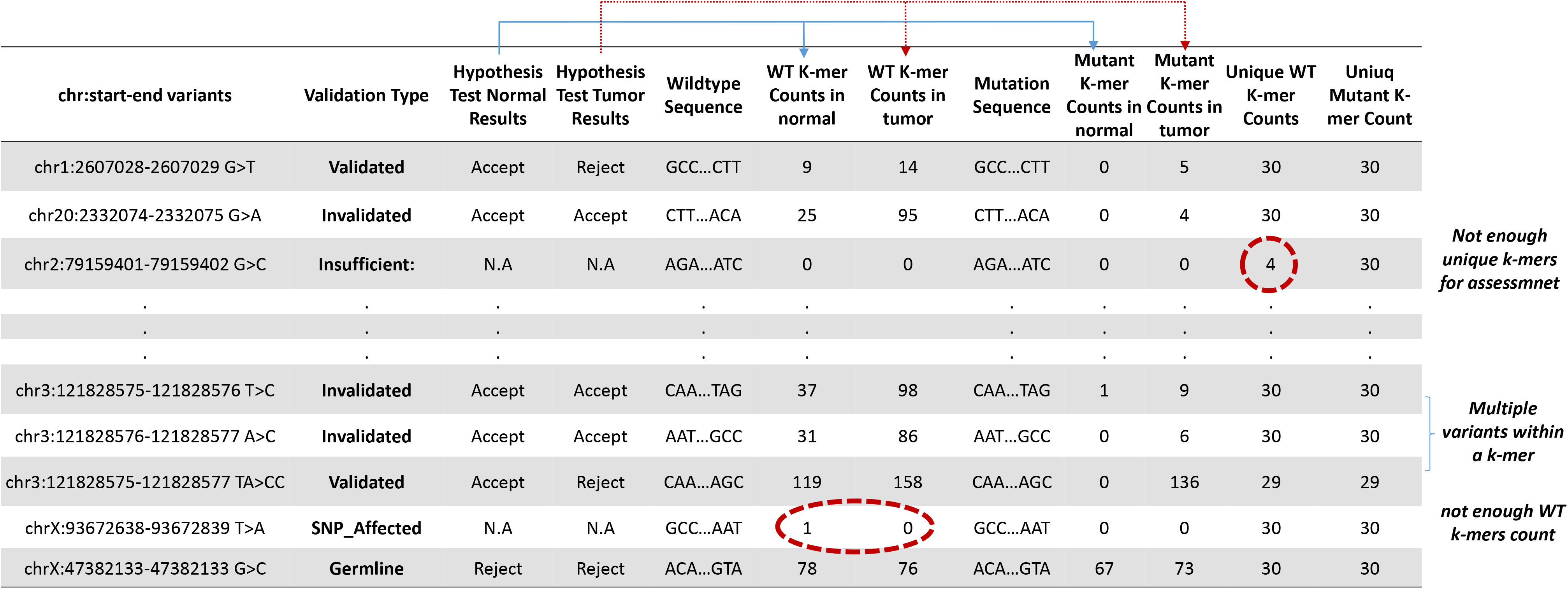
Final summary table. The final summary table with the validation status of variants by KmerVC.

We consolidated the software’s analysis and results into one final summary **(Figure 2)**. In the final summary table, we organize the k-mer counts by mutations and determine if the counts of mutant k-mers sufficiently verify the mutations using binomial tests that assume a sequencing error rate of 1%. The summary table has reporting headers and 17 columns **(Supplementary Table S1)** with information regarding the variants and their surrounding k-mer regions.

### Reference and sequence data

#### Reference genome and annotations

We downloaded GRCh38 from the National Center for Biotechnology Information (www.ncbi.nlm.nih.gov). Only the canonical chromosomes and chromosome M were used in the analysis. The coordinates of gaps (N) and repeats were obtained from UCSC genome browser; gap.txt.gz. For the definition of coding sequence, we downloaded Consensus Coding Sequence (CCDS) from NCBI.

#### Simulated data

We simulated a sequence data set containing substitutions, insertions, deletions and indels. A segment of human GRCh38 chromosome 1 from positions 629640 to 5629640 (50 Mb) was used as the reference sequence for this in silico data. Then, we introduced mutations at 120 random positions into this segment. The indels had six different base pair lengths; 1, 3, 5, 10, 15, and 20. Two different data sets were generated with one representing ‘normal’ and the other containing the mutations. We used the read simulator wgsim (http://github.com/lh3/wgsim) to generate simulated short paired-end reads with 100 bp length at a depth of 25 (normal) and 50X coverage (mutation-containing). This simulation incorporated a 1% sequencing error model. The following command was used:

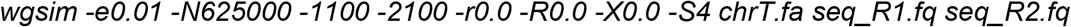

The simulated sequencing data is publicly available at our website as listed below: (https://dna-discovery.stanford.edu/publicmaterial/software/kmervc/simulation/).

#### Exome sequencing data from TCGA

We downloaded exome sequencing data (BAM files) of 50 colorectal cancers from the Genomic Data Commons (GDC) data portal (portal.gdc.cancer.gov). FASTQ files were derived from BAM files using Picard SamToFastq (version 2.9.0). We also downloaded four Mutation Annotation Format (MAF) files generated by MuTect2, SomaticSniper, MuSE, VarScan2 Variant Aggregation and Masking from the GDC data portal; i) TCGA.COAD.mutect.03652df4-6090-4f5a-a2ff-ee28a37f9301.DR-10.0.somatic.maf.gz, ii) TCGA.COAD.somaticsniper.70835251-ddd5-4c0d-968e-1791bf6379f6.DR-10.0.somatic.maf.gz, iii) TCGA.COAD.muse.70cb1255-ec99-4c08-b482-415f8375be3f.DR-10.0.somatic.maf.gz, iv) TCGA.COAD.varscan.8177ce4f-02d8-4d75-a0d6-1c5450ee08b0.DR-10.0.somatic.maf.gz. We only included variants with “PASS” in the “FILTER”.

#### In silico sequence data for insertions and deletions

To test the performance of our approach for validating indels, we examined a category of sequence which is one of the most challenging cases. Specifically, microsatellites are composed of simple tandem repeats **(STRs)** which are present throughout the human genome. STRs have different classes of repeat motifs that include mono-, di-, tri- and tetranucleotide sequences. Microsatellites are prone to accumulating indel mutations at a high frequency and are extremely difficult to identify reliably given their repetitive structure.

We generated virtual VCF files that could be used as inputs. First, we located microsatellites by searching for particular mononucleotide repeat motifs. This process involved matching of single nucleotide repeats against the reference genome. This method gave us a total of 5,751 microsatellite regions. Next, we identified insertions or deletions of up to three bases occurring in these microsatellite regions. Therefore, six variant calls were generated: three deletions and three insertions with each of the microsatellite’s repeated nucleotide with lengths from one to three base pairs. Finally, we created VCF files recoding the simulated insertions and deletions at these regions and utilized these files as input to the KmerVC to evaluate their significance as plausible mutations. This file, MS_DNA_indels_grch38.vcf, is available at https://github.com/compbio/kmerVC.

## RESULTS

### Overview of k-mer evaluation

Our analysis of k-mer counts relied on the analysis of matched samples to confirm candidate mutations **(Figure 3)**. The use of matched normal versus tumor sample enable us to eliminate germline polymorphisms, rare variant and hereditary mutations. We determined the properties of a somatic mutation and how it generates a series of novel (neo) k-mers.

**Figure 3.**
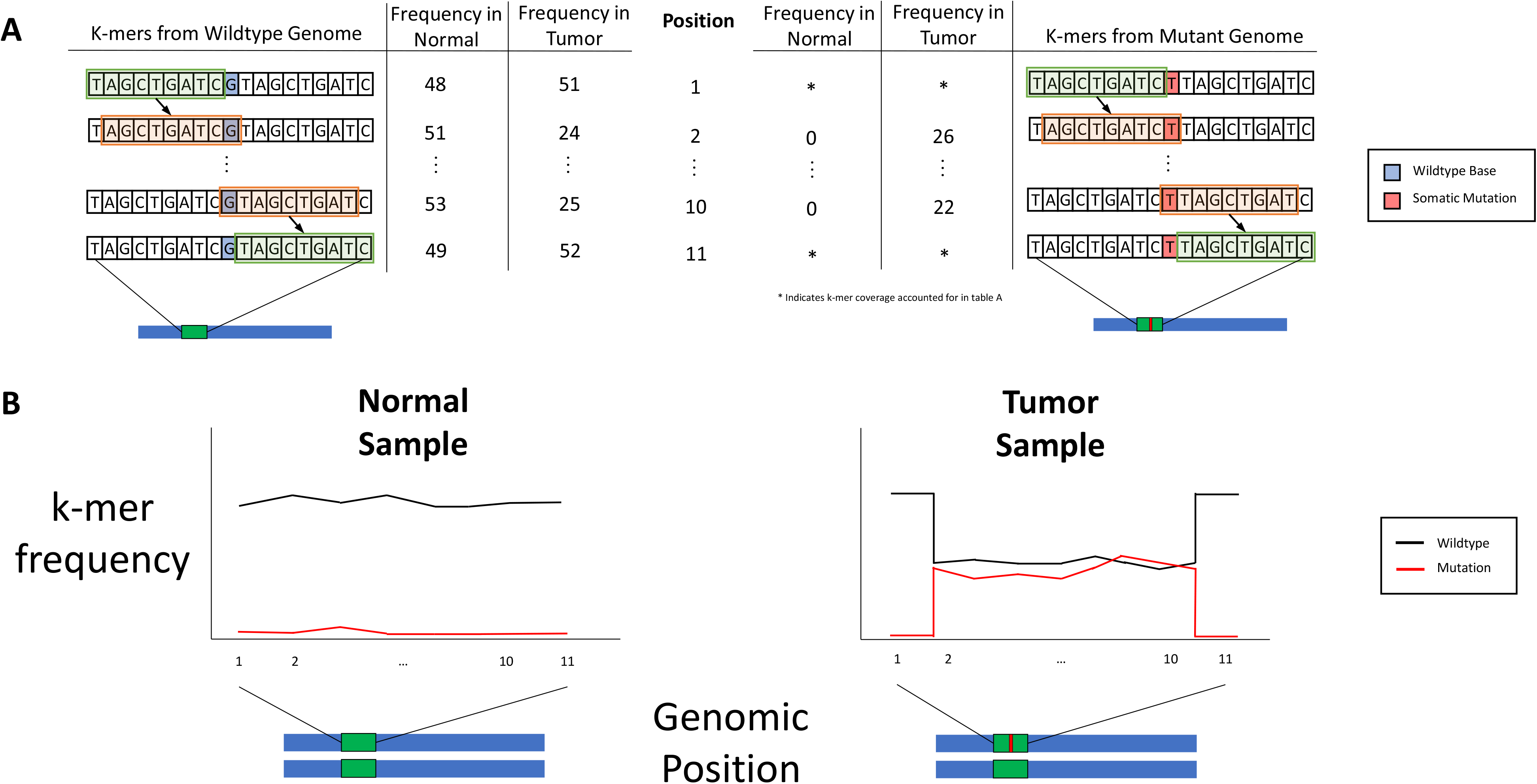
Principle of k-mer counts for variant detection. (**a**) The majority of mutations generate neo k-mers. **(b)** The counts of k-mers are affected by mutations. The mutation is noted by a red bar occurring with an exon which is denoted by a green box.

These sequences are different than k-mers present in the reference genome or the matched germline comparison. These neo k-mers identify the somatic mutations in the tumor. For instance, a substitution such as G to T will generate nine neo K-mers (length of nine bases) as seen in **Figure 3A**. We utilized a sliding window to obtain the k-mers of the designated lengths spanning the mutation region of interest; this provided both mutation-containing and wild-type k-mers. A normal genome yields no mutation k-mers while a tumor sample with mutations yields a significant number of mutant k-mers. Similarly, the number of corresponding wild-type k-mers in the tumor will be lower if the mutation is heterozygous and diminished if the mutation is homozygous. For example, with a coverage of 50X the count of mutant k-mers is expected to be 50 for homozygous and 25 for heterozygous mutations in the tumor sample, assuming a 100% tumor purity **(Figure 3B** illustrates a heterozygous example). A difference in the counts of wild-type and mutant k-mers between the matched normal and tumor sequencing data allows for the statistical assessment of the validity of a somatic mutation variant.

### Evaluation of genome-wide base positions by unique k-mers

For this study, one of the most important properties of a k-mer is the uniqueness of its sequence compared to the total number of k-mers present in the human genome reference (GRCh38). Specifically, this means whether a given k-mer appears only once in either the forward or reverse direction of the genome reference. We postulated that mutations appearing in these unique “neo’ k-mers have specific properties that make them readily distinguishable **(Supplementary Figure 1)**.

We determined the number of unique k-mers in the reference. All k-mers in the reference genome were counted using one bp increments over its length. When the length of a k-mer is greater than or equal to 19 bases, more than 90% of distinctive k-mers are uniquely represented in the human genome reference **(Supplementary Figure S2)**. When k-mers are 30 bases in length, 96.2% of these sequences are unique. Based on our analysis, k-mer lengths of greater than 30 bases lead to only minimal increases (less than 2%) in the proportion of unique k-mers among distinctive k-mers **(Supplementary Figure S2)**. This result has been confirmed elsewhere (22).

It is important to note that the length of k-mers directly determines the total fraction of unique k-mers. However, we found that increasing the size of k-mers also lead to greater probability of sequencing error artifact. Specifically, we observed that there was a tradeoff in that the distribution of longer k-mers and their properties could be artifactually skewed by sequencing errors. Shorter length k-mers were less prone to artifact from these errors. Another study confirmed this issue (18). Thus, our software allows users to select different k-mer lengths for their studies.

Having identified 30-mers as a length with suitable properties, we identified the total number of detectable bases within the human genome reference. We used the following definition of unique k-mer as related to individual base positions in the reference. If one examines any given position in the human genome reference, there are a total of 30 k-mers (length = 30) which overlap this base. Stating it differently, a sliding window provides a series of k-mers that cover a given base position **B** with the first having **B** as its last position and the last having **B** as its first position. So long as one of these 30-mers is unique, this base is considered mappable. For our analysis, we defined several terms. We defined a ‘**detectable base**’ as one which overlaps five or more unique 30-mers. This metric provides robust and unique mapping from multiple 30-mers even when they contain regions with sequencing errors.

Similarly, we conducted an extensive analysis of genome positions which are not mapped with unique k-mers. We used the term ‘**dark base’** to refer to a base position that is covered by less than five unique 30-mer **(Figure 4A)**. We used the term ‘**dark region’** to refer to a segment of sequence containing two or more adjacent dark bases. These are segments of genome sequence that may have issues aligning correctly to the reference. We provide the coordinates of dark regions as BED files (dark_bed.zip), which can be downloaded from https://dna-discovery.stanford.edu/publicmaterial/software/kmervc/.

**Figure 4.**
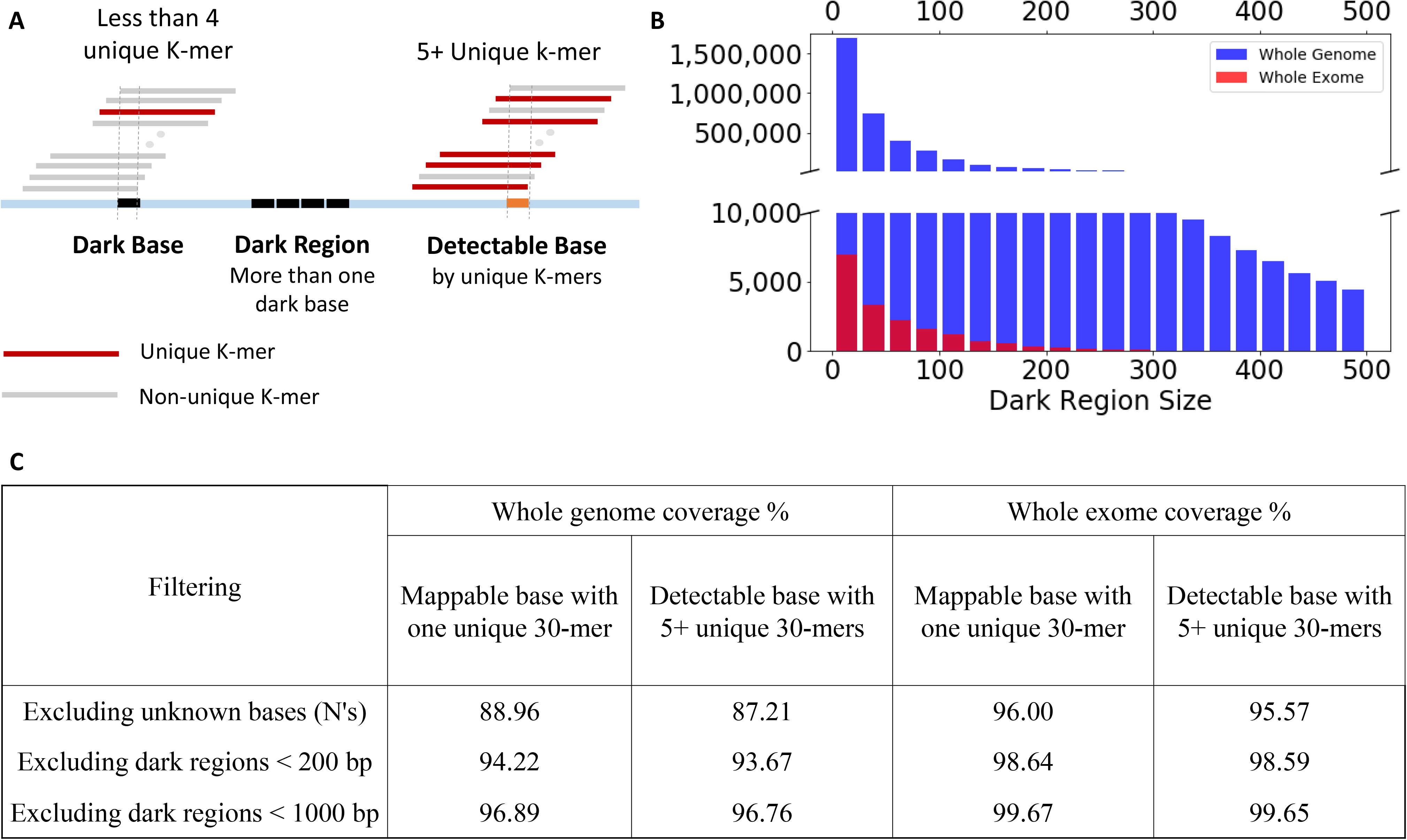
The characteristics of detectable and dark. **(a)** Definition of detectable, dark base / region. **(b)** The distribution of size of dark regions. **(c)** Coverage of unique k-mers in GRCh38 after excluding dark regions.

Based on these definitions, we determined that 86% of the reference genome bases can be mapped using unique k-mers. As an added filter, we eliminated the unknown bases (typically annotated with the character *N*), which comprise 5.3% of the reference genome and make up a total of 164Mb. As a result, we found that 87.2% of the known bases in genome reference are detectable by at least five or more unique 30-mers. This high level of overall coverage based on unique k-mers is remarkable given that more than two-thirds of the human genome is composed of repeat elements and low complexity regions (23).

If one considers only exon regions, which total ~33Mb according to CCDS (24), 95.6% of the exon bases are detectable by at least five or more unique 30-mers. The exons of established cancer genes such as *TP53*, *APC* and *KRAS* have a very high portion (94.6%) of detectable bases on median **(Supplementary Table S2)**.

We determined that these dark regions (i.e. gaps in mappable bases) generally were less than 150 bases in length **(Figure 4B)**; this length is shorter than the typical sequence read from an Illumina system. We identified adjacent unique 30-mers as either upstream or downstream anchors of the dark region within the length of a single sequence read (i.e. 150-300 bases) or within the insert size distribution that occurs for paired end reads (i.e. 150 – 500 bases). With this extended definition of detectable base using pair-end reads, we found that the detectable regions of whole genomes and exomes increased from 87.21% and 95.57% to 93.67% and 98.59% respectively **(Figure 4C)**. With this observed increase in exon detectable bases, there were no dark regions found in 29,588 (88.6%) of the 33,381 genes annotated per CCDS. If one considers the long sequence reads available from Oxford Nanopore or Pacific Biosciences, these dark regions can be further reduced in size and more bases can be identified using our k-mer approach. This prediction can be seen in the theoretical example of sequence read lengths of 1Kb **(Figure 4C)**.

There are other resources that consider the “mappability” of k-mers. For example, the Umap resource available from the UCSC Genome Browser provides mapping information for sequences of varying lengths, ranging from 24 to 100 bases (25). This feature is used for determining which sequences can be accurately aligned to the reference genome using k-mers. Our method is different in several ways. First, we define a single base as “detectable” if it overlaps with five or more unique k-mers. Using Umap’s formula, our “detectable” base has the minimum “mappability” of 0.167 (=5/30). We use this definition of a “detectable” base to validate somatic mutations which has not been described for Umap. Our method also allows us to introduce and characterize the “dark regions” of the genome which had previously been considered as being essentially unmappable.

### Validation of simulated variants from in silico sequence data

We evaluated the feasibility of k-mer counts to validate the identification of a ground truth set of mutations generated in silico. We applied the KmerVC program to a simulated sequencing dataset of a wild type genome with three simulated sequencing data files generated from the mutated genome that contain; i) 120 substitutions, ii) 120 insertions, and iii) 120 deletions respectively **(see Methods)**. The precision and recall values were calculated with the following formula; Precision = TP / (TP + FP) and Recall = TP / (TP + FN) where TP is True Positive, FP is False Positive, and FN is False Negative. TP indicates the number of correctly validated simulated mutations. FP indicates the number of validated mutations that were not one of the simulated ones. FN indicates the number of simulated mutations that were not validated.

We generated a ground truth set of mutations embedded within in silico sequence data. Categories of different mutations included: i) 120 simulated substitutions + 380 random substitutions + 1,352 raw variants from GATK 3.8; ii) 120 simulated insertions + 380 random insertions + 1,346 raw variants from GATK 3.8.; iii) 120 simulated deletions + 380 random deletions + 1,362 putative variants from GATK 3.8. Substitution mutations were provided to the program as a BED file. We provided the actual sequences with the indel mutations seeing that micro-homology motifs complicate the use of coordinates for reporting the location of either insertions or deletions.

**Figure 5** shows the precision and recall of KmerVC for simulations based on an alpha value of 0.01 with multiple test correction. We tested different lengths of k-mers and found that 30-mers performed better than 20-mers, but there was no significant improvement with 40 or 50-mers **(Supplementary Figure S3)**.

**Figure 5.**
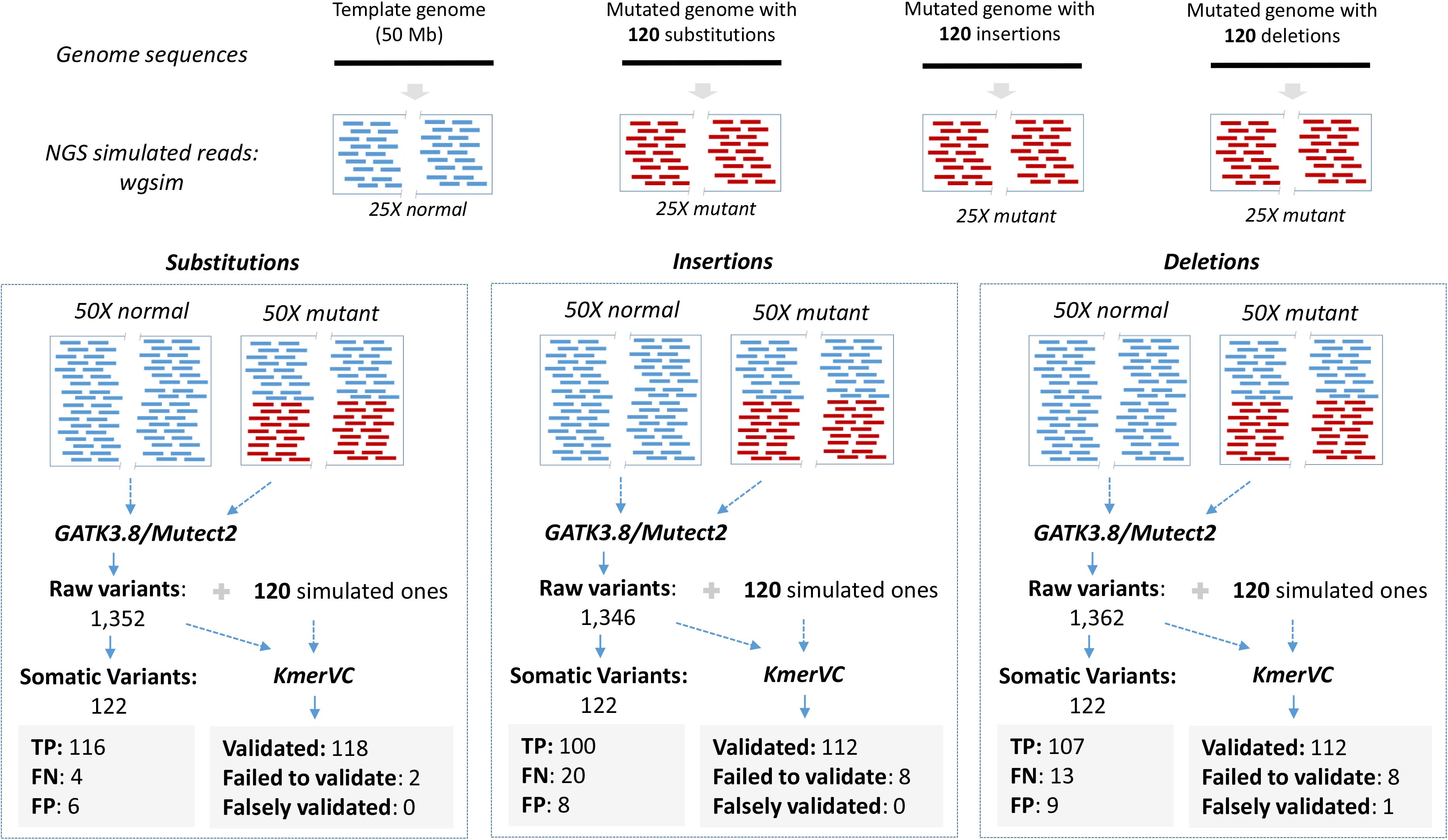
Validation of simulated substitutions, insertion and deletions in sequencing data.

From the ground truth sequence data, we used Mutect2 to call mutations. This program was run in both its GATK 3.8 and GATK 4 versions. Because the upgrades of GATK 4 involved improvements in specificity of variant calling, we observed a significantly lower performance in detecting indels in the simulated data. This calling decrement was apparent even after extensive filtering. Therefore, we used variant calls from GATK 3.8. The performance of the Mutect2 pipeline is shown in **Figure 5**. When we analyzed the GATK substitution variants, Mutect2 called 116 TPs, 4 FNs, and had 6 FPs. Of the four FNs, one was not called at all while three did not pass the clustered_events filter (i.e., proximal variant calls failed on the assumption that the clustering is a proxy for mapping artifacts). Against the insertion data, Mutect2 found 100 TPs, 20 FNs, and had 8 FPs. Two of the FNs were not called at all; 7 failed at least in part because of the clustered_events filter, while 10 failed because they failed to pass the triallelic_sites filter. This situation occurs when at least two candidate alternate alleles are found to be equally likely; since this is unlikely to occur in a tumor, it is assumed to be evidence of an artifact. Against the deletion data, Mutect2 found 107 TPs, 13 FNs, and had 9 FPs. Here, five true positives were not called, four were filtered by the clustered events filter, and two were filtered because they involved deletions of tandem repeats. As we noted previously, Mutect2 had had reduced performance for calling insertions with a lower recall rate relative to deletions.

Next, we ran KmerVC to validate the simulated variants along with all raw the Mutect2 calls. Our program confirmed nearly all of the ground truth mutations successfully and invalidated nearly all other tested mutations regardless of variant types **(Figure 5)**. Remarkably, all FPs but one, which called by Mutect2, were not validated by KmerVC and thus invalidated them properly. Furthermore, KmerVC successfully validated 2 out of 4 FNs in substitutions, 17 out of 20 FNs in insertion and 5 out of 13 FNs in deletions; our program was able to increase the number of TP from FNs. Overall, these results indicated that our k-mer analysis validated true positives while avoiding false calls made by a well-established variant caller.

### KmerVC analysis of TCGA exome data

Next, we determined our application’s performance for validating mutations from actual exome sequence data representing 50 normal tumor pairs. We downloaded the matched normal/tumor FASTQ files and VCF files derived from four pipelines (Mutect2, Varscan, Muse, and somaticsniper), all available from the TCGA data portal. The VCF’s for each mutation caller are available separately for each caller pipeline for the 50 cancer samples and separated into two groups: i) substitution and ii) indels. For example, Mutect2 called a total of 21,601 substitutions and 3,412 indels across the 50 tumor samples.

#### Substitutions

We used KmerVC to process the FASTQ and VCF files of each sample. For our first analysis, we examined the Mutect2 calls. Our analysis determined which of 21,601 Mutect2 substitutions could be validated for each sample. For any given normal tumor pair, KmerVC successfully validated on average 91% of Mutect2 variants for 0.01 alpha value and 83.2% after multiple test correction **(Supplementary Table S3)**. The difference in the percentages that were validated is an indicator of the smaller alpha value due to Bonferroni correction; a lower alpha value imposes a more stringent threshold which excludes lower quality variants.

To determine what features distinguished the validated versus non-validated mutations, we conducted a manual inspection of the calls from Mutect2 that were not validated by KmerVC in the tumor sample of TCGA-AA-3350. KmerVC failed to validate four out of 111 mutations derived from Mutect2. Three of the four missed mutations were due to proximal SNPs while one was due to the mutation being located at the end of most of the reads.

With alpha value of 0.01 with multiple test correction, KmerVC validated 89%, 91%, and 94% of variants derived from Varscan, muse, and somaticsniper respectively on average **(Supplementary Table S3)**. Interestingly, the number of Mutect2 variant calls validated by KmerVC was larger than the number of Mutect2 variants also identified by any of the three other variant callers **(Supplementary Table S4)**.

#### Insertions and deletions

We analyzed indels among these 50 tumors. Our validation analysis required there to be a flanking base on both sides of the mutation to ensure the length of insertion and the indel’s identity. Therefore, the number of affected k-mers was extrapolated using the size of the indels. For instance, a two-base insertion in turn overlaps with 27 of the surrounding 30-mers.

We examined all 3,412 indels identified in 50 samples. On average, KmerVC validated the greater majority of indels (75%) derived from Mutect2 with a corrected alpha value of 0.01. The percentage of validated indels went down to 71.7% if one decreased alpha value by Bonferroni correction **(Supplementary Table S5)**. In conclusion, KmerVC validated most substitution calls as well as indels identified by Mutect2 with a high performance. The KmerVC outcomes (tcga.zip) for all TCGA samples are available at https://dna-discovery.stanford.edu/publicmaterial/software/kmervc/

### Validating microsatellite mutation

KmerVC has the flexibility to validate different classes of mutations. One of the most challenging somatic mutations to identify are indels present within microsatellite **(MS)** sequences. To determine if our approach had applicability to verifying MS indels, we generated a VCF file that contains potential indels at 5,751 MS DNA in the coding regions **(Methods)**. KmerVC successfully validated 29.5 (median) indels at MS DNA per sample ranging from 4 to 574 MS DNA with corrected alpha value of 0.01 (**Supplementary Table S6)**. We examined sequence alignment plots by Golden Helix GenomeBrowse 3.0.0 (www.goldenhelix.com), and visually confirmed the presence of a set of these indels **(Supplementary FigureS4)**.

## DISCUSSION

Our study examined the properties of k-mers from the human genome reference. With this information, we developed an application that validates somatic mutation calls for true positives using the characteristics of counts and uniqueness extrapolated from primary sequencing data. We note that our approach provides a way of rapidly validating indels. One can imagine strategies where the indel threshold can be lowered to improve sensitivity with follow up KmerVC analysis and validation to improve specificity.

We evaluated the performance of KmerVC using simulated sequencing data and exome data from TCGA. KmerVC achieved high performance by validating more than 93% of true mutations with nearly zero falsely validation based on the simulation data for all variant types: substitutions, insertions, and deletions. Furthermore, KmerVC successfully validated most of the variants derived from Mutect2 in TCGA exome sequencing data. Remarkably, we observed that KmerVC rarely validates any variants falsely, which is a strength in precision medicine.

Several studies have utilized the k-mer count for identifying previously known germline mutations or indels. However, there are few if any studies examining the application of k-mer counts to evaluating somatic mutations in cancer sequencing data. As demonstrated in this study, KmerVC validates the mutation of interest with a straightforward binomial test using only four numbers; i) wild-type counts in normal, ii) mutation counts in tumor, iii) wild-type counts in tumor, and iv) mutation counts in tumor. The simplicity of this metric enables us to validate variants easily without the need for hidden heuristics or complex optimizations. This feature is extremely useful for identifying cancer mutations of interest when sequencing data is available from multiple different projects. The counts of expected k-mers for these mutations can be quickly compared between samples. Any validated variant calls by KmerVC can be compared among different samples easily, providing an added level of quality control.

Our method has some limitations. First, insert sizes larger than the k-mer length cannot be detected. For a future study, we plan to extend our approach to detect structural variations, which should generate novel combinations of k-mers. These potential features may allow us to identify insertions larger than the k-mer size. Second, some regions do not overlap with any unique k-mers despite the high number of unique k-mers across the human genome reference. Our characterization of the dark regions of the genome highlights our use of unique k-mers. Moreover, different lengths of sequence reads that cover these mutations provides a number of opportunities to improve validation analysis. Third, KmerVC is strictly a validation program of variant calls and does not account for false negatives. These unaccounted variants may arise from mutation-related k-mers that are not unique or expected k-mer counts for a given coordinate position that occur as the result of a single nucleotide polymorphisms **(SNPs)**. The use of a matched normal samples provides way of eliminating SNPs. However, we anticipate that a substantial number of somatic mutations lead to neo k-mers which are unique and this property may enable us to identify false negatives and provide a way to transition our tool into a full-fledged mutation caller.

In conclusion, for this study, we comprehensively examined the landscape of mappable regions by unique k-mers in the human genome. Using the characteristics of k-mers, including their uniqueness in the reference and the ease of counting them, provides an excellent way to determine if called mutations are true positives, especially for indels.

## AVAILABILITY

The kmerVC program and all simulated sequencing data are available in the GitHub repository (https://github.com/compbio/kmerVC). All whole exome sequencing data are available at TCGA data portal (https://portal.gdc.cancer.gov/).

## ACCESSION NUMBERS

Not applicable.

## SUPPLEMENTARY DATA

Supplementary Data are available at NAR online.

## ACKNOWLEDGEMENTS

The work is supported by the National Institutes of Health [2R01HG006137-04 to H.P.J., P01HG00205ESH to J.M.B AND H.P.J., U01HG010963 to H.J.L., D.P. and H.P.J.].

Additional support for HPJ came from the Research Scholar Grant, RSG-13-297-01-TBG from the American Cancer Society, National Science Foundation Award 1807371 and the Clayville Foundation. We wish to thank Stephanie Greer, Billy Lau, and Sue Grimes for discussions and comments on the manuscript.

## AUTHOR CONTRIBUTIONS

HJ.L. and H.P.J. conceived and designed the study; A.S., HJ.L., and D.P. implemented the computational method. HJ.L and D.P. developed the statistics. A.S. wrote the Python script.

J.M.B analyzed simulation data using the different versions of GATK. HJ.L. and H.P.J. analyzed the data. All authors contributed to the manuscript writing. HJ.L and H.P.J. supervised the project.

