## Supplementary Figures for "Unique K-mer sequences for validating cancer-related substitution, insertion and deletion mutations": Supplementary_Data.pdf

Supplmentary Materials includes 4 figures and 6 tables.

**Supplementary Figure S1**.

A

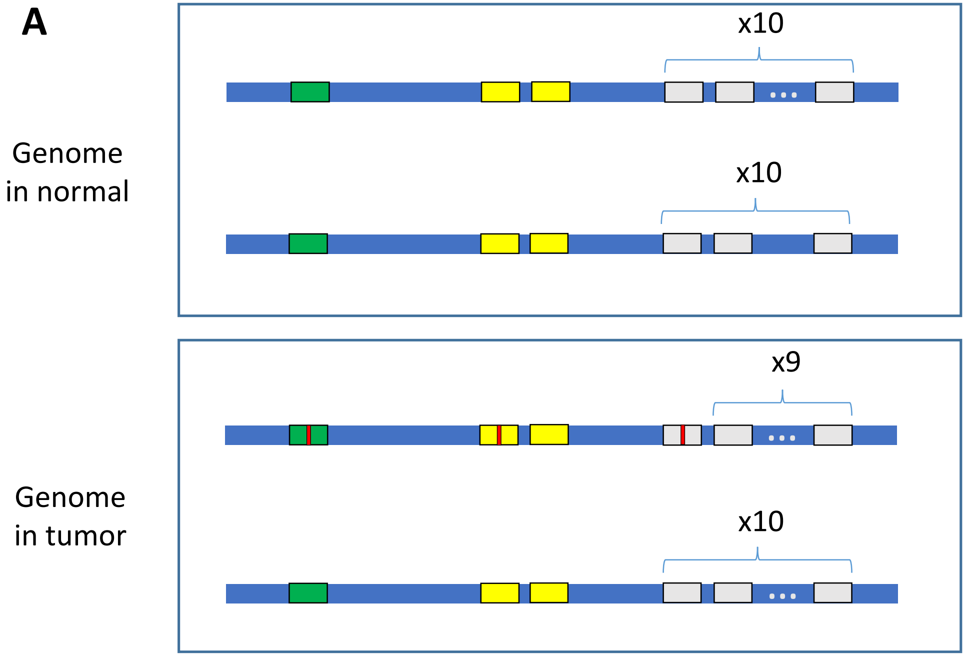


Expected counts of k-mers from 50X coverage with 100% tumor purity

| K-mer type | Wildtype k-mer counts | | Neo k-mer counts | | % of neo k-mers |
|---|---|---|---|---|---|
| | Normal | Tumor | Normal | Tumor | |
| | 50 | 25 | 0 | 25 | 50 |
| | 100 | 75 | 0 | 25 | 20 |
| | 500 | 475 | 0 | 25 | 5 |

**Supplementary Figure S1**. The strong signal from unique k-mers. (A) depicts the genomic strands of the patient's normal and tumor samples with particular existing green, yellow, and grey reads outlined that occur once, twice, and ten times respectively. We depict a somatic mutation on one green, yellow, and grey read in the tumor sample. Noticeably, (B) shows that the percentage of neo-kmers decreases as we deal with non-unique reads and k-mers that exist in greater frequency. The unique k-mers shows the mutation events clearly.

**Supplementary Figure S2**

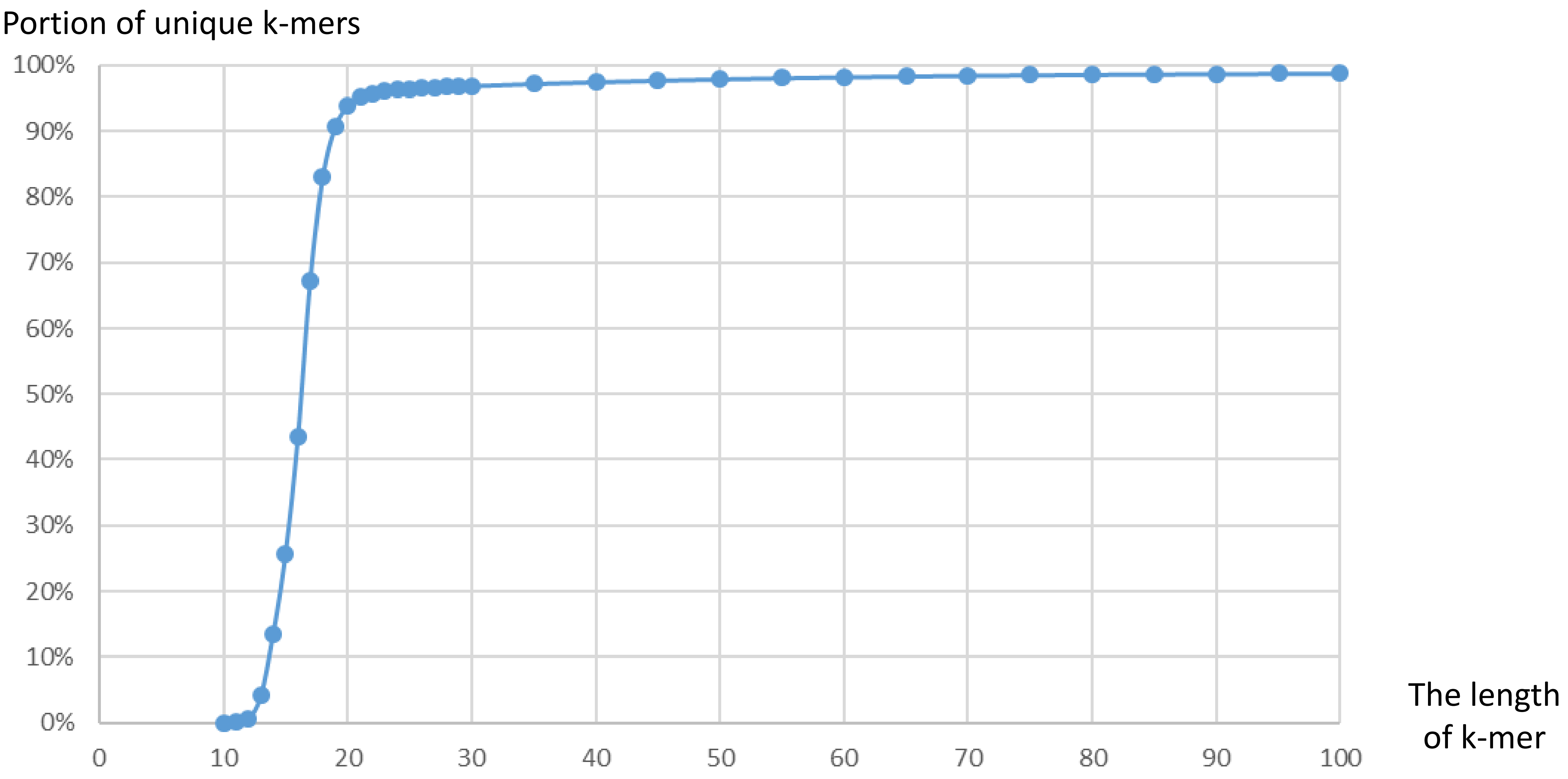


**Supplementary Figure S2**. Percent of unique k-mers out of all distinctive k-mer by k-mer length.

**Supplementary Figure S3**

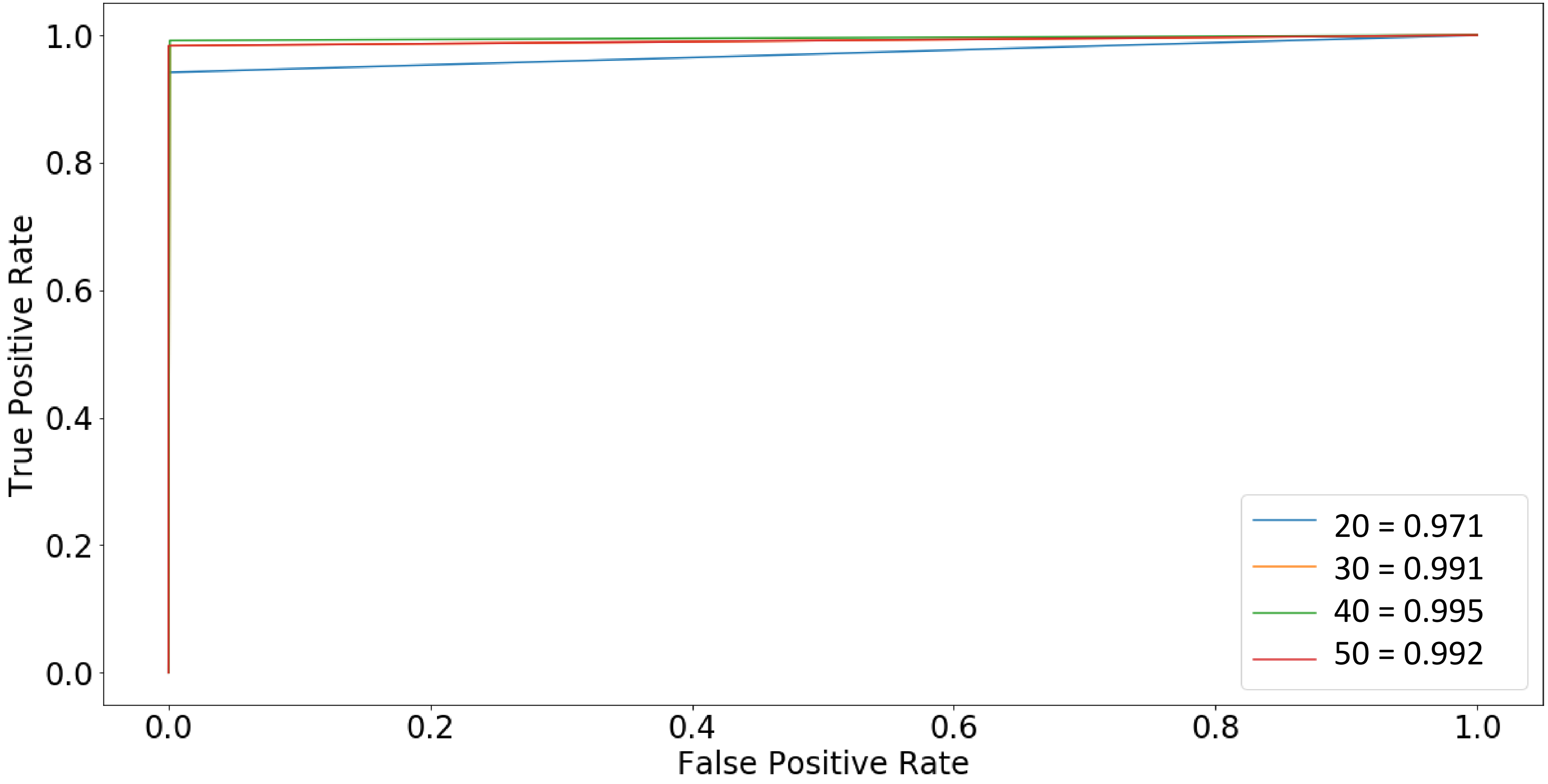


**Supplementary Figure S3**. AUC curves by different length of k-mers

### Supplementary Figure S4

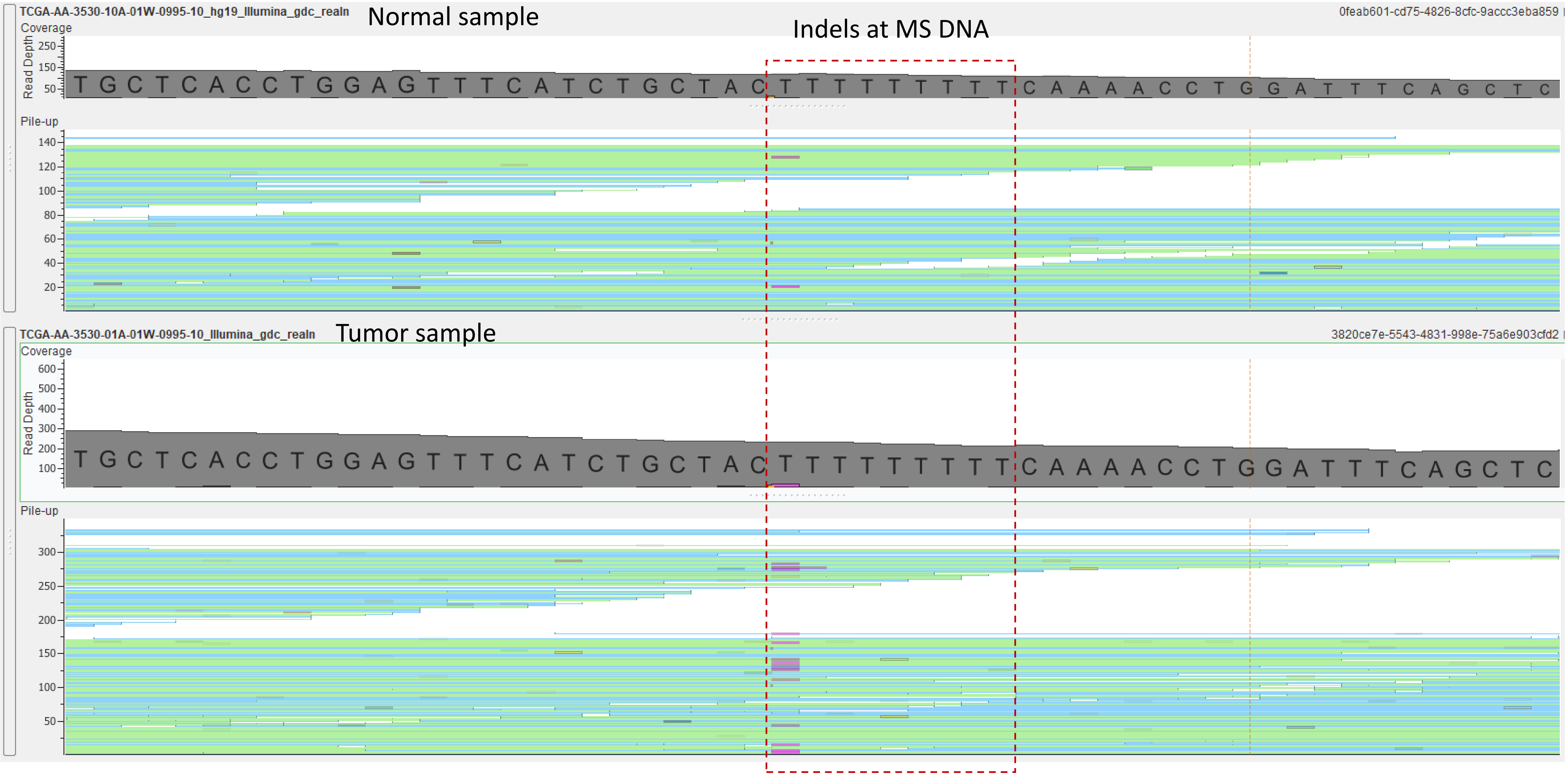


**Supplementary Figure S4**. (A) One T deletion at Chr1: 12,725,526

#### Supplementary Figure S4

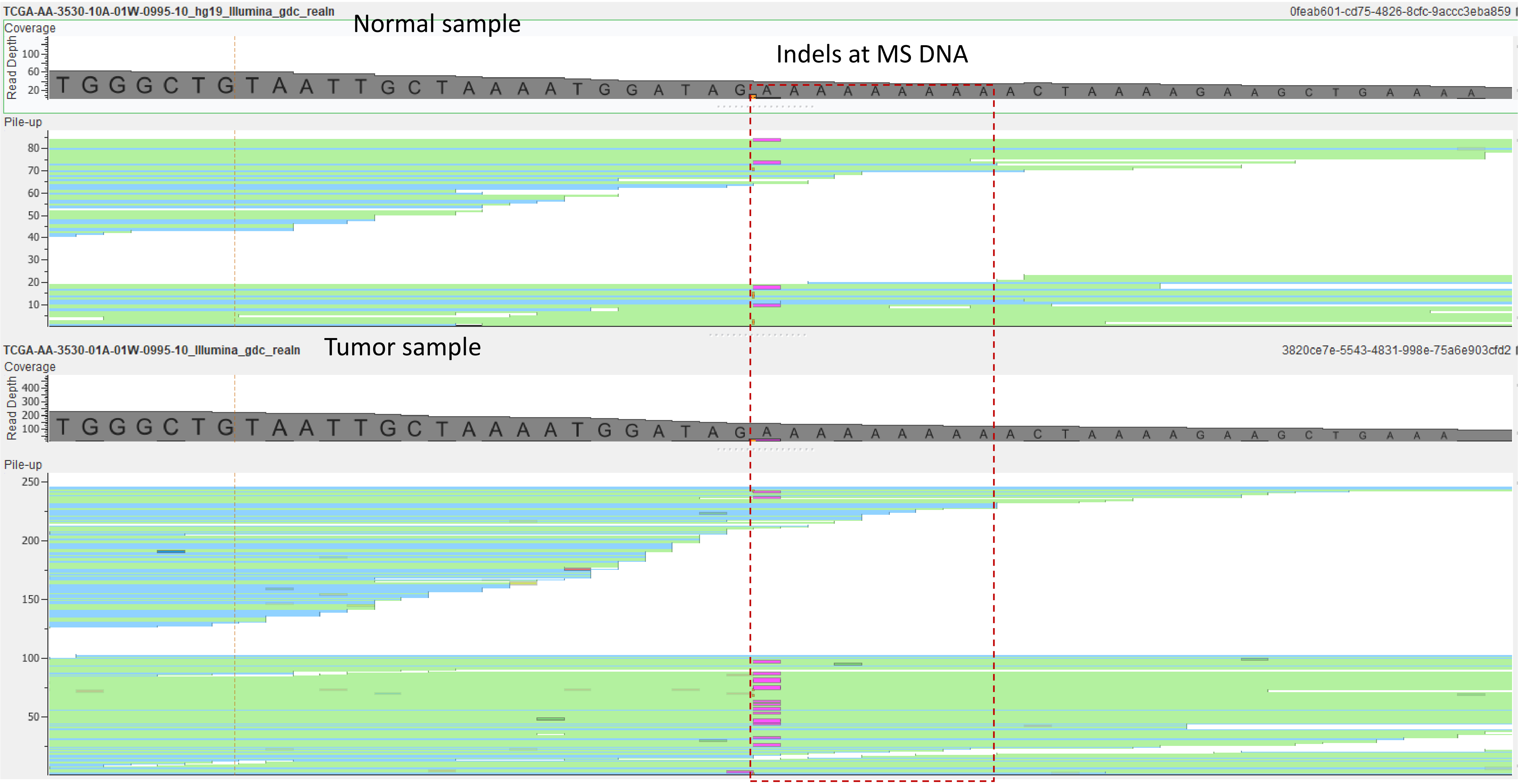


**Supplementary Figure S4**. (B) One A deletion at Chr1: 114994980

### Supplementary Figure S4

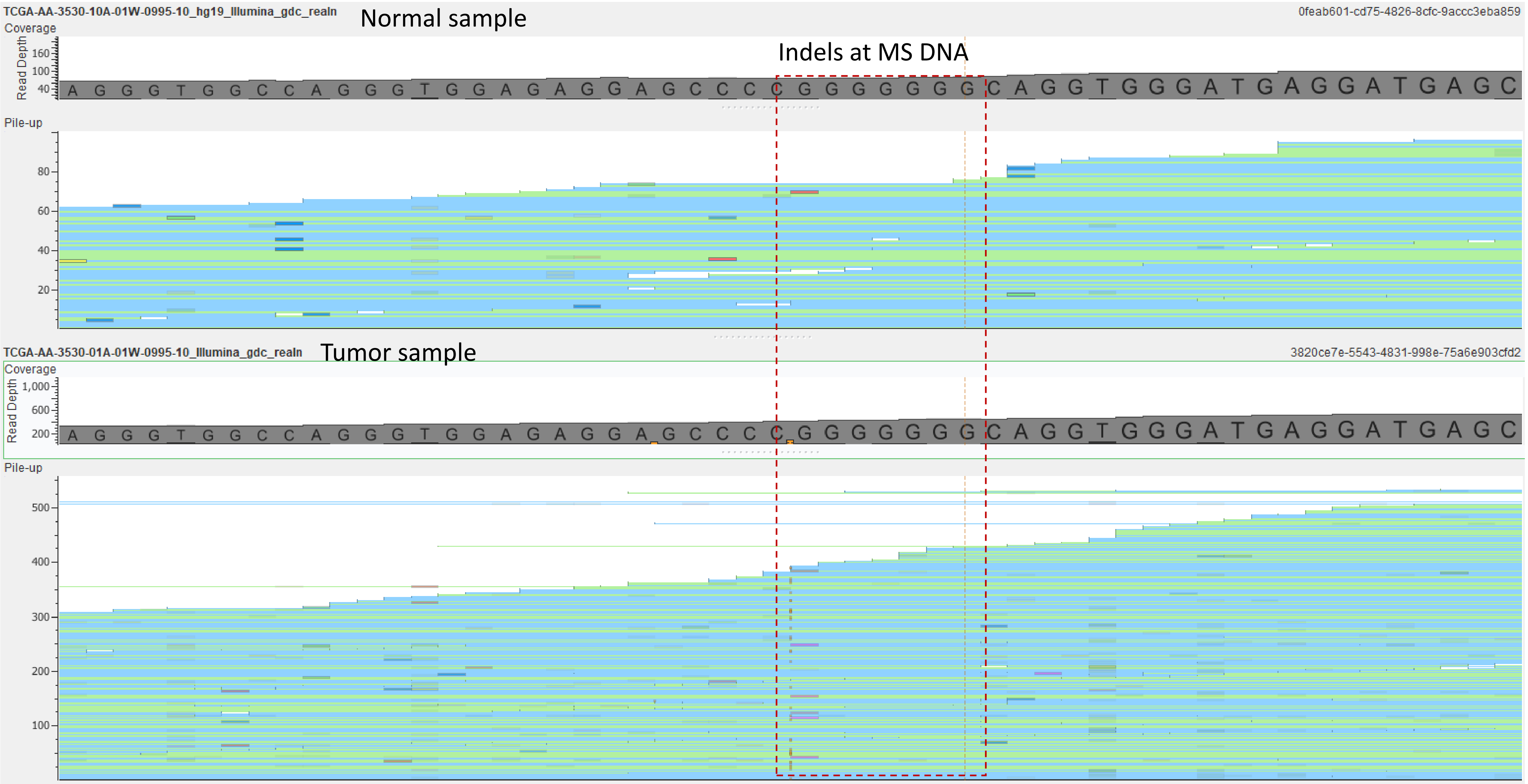


**Supplementary Figure S4**. (C) One F insertion at Chr20:4182380

#### Supplementary Figure S4

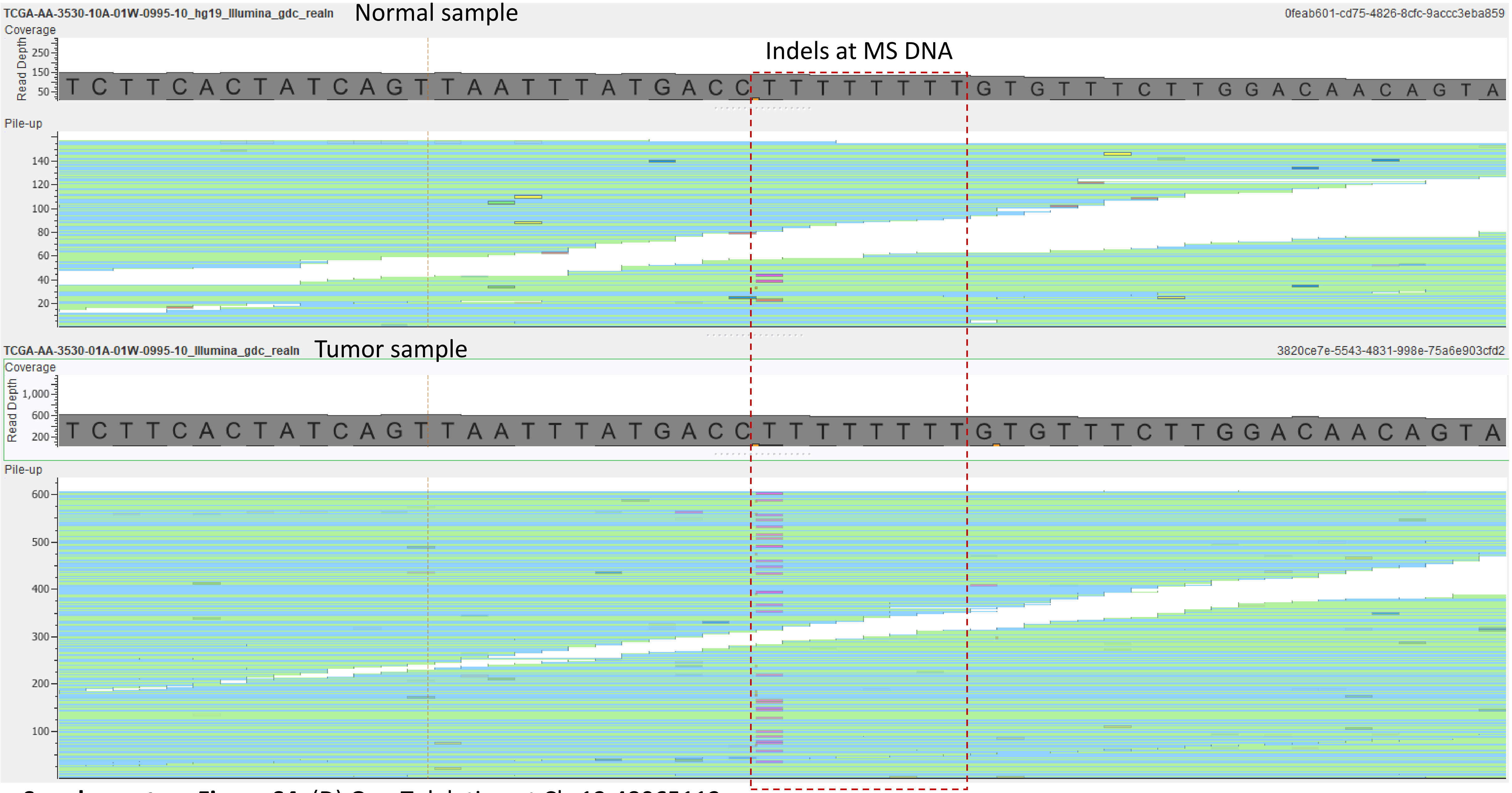


**Supplementary Figure S4**. (D) One T deletion at Chr12:48065113

### Supplementary Figure S4

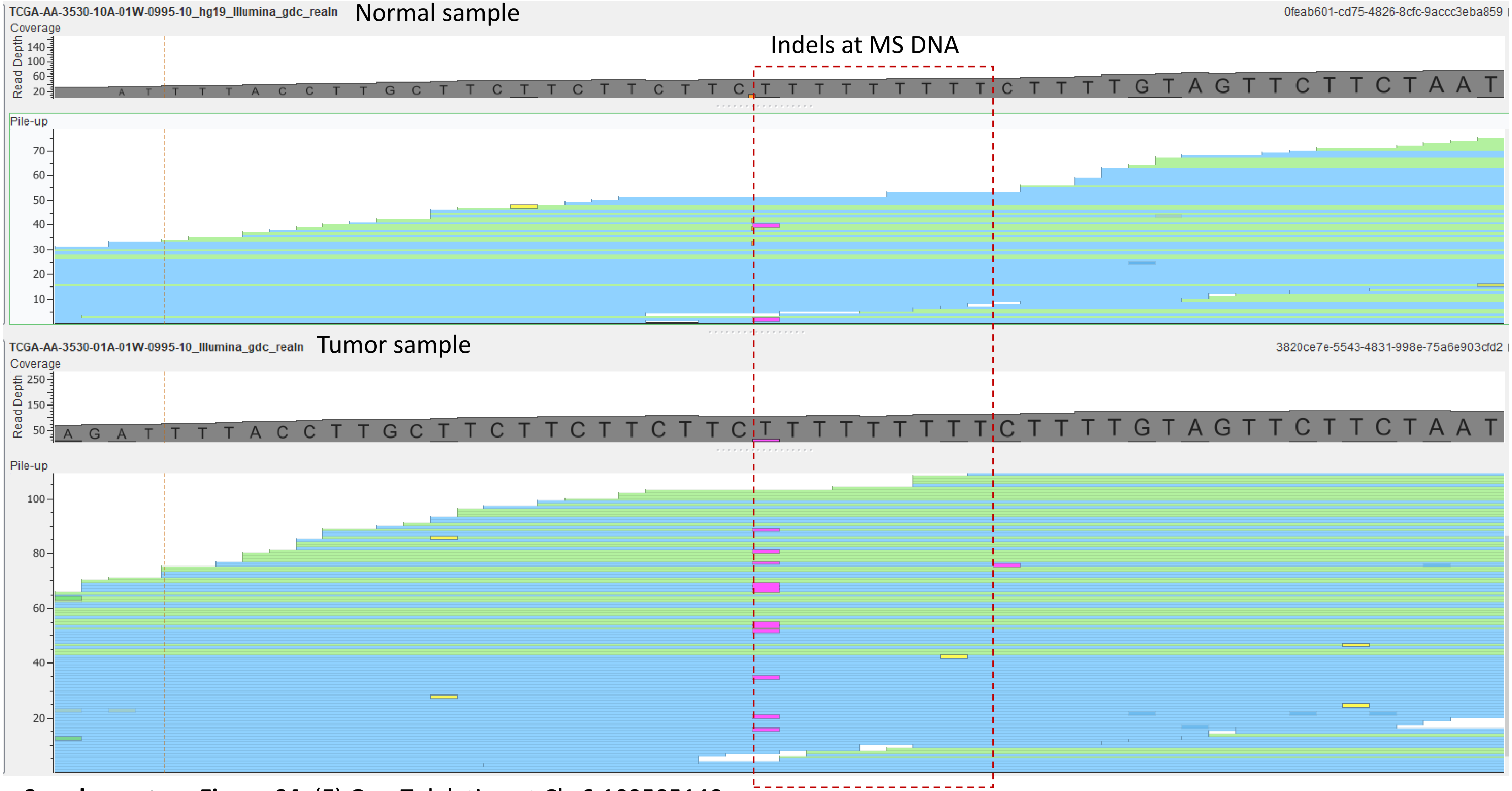


**Supplementary Figure S4**. (E) One T deletion at Chr6:109585140

#### Supplementary Figure S4

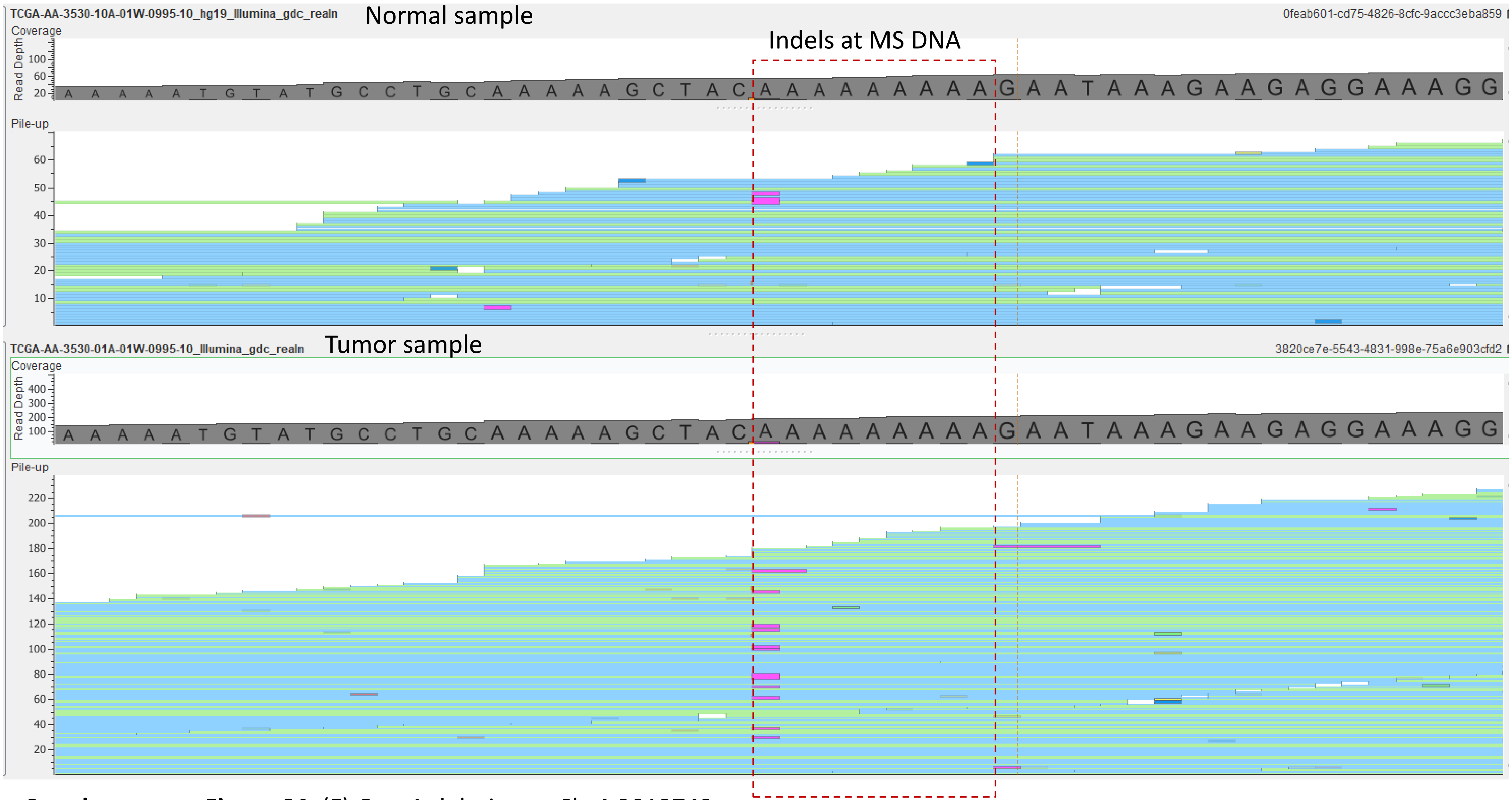


**Supplementary Figure S4**. (F) One A deletion at Chr4:3013743

#### SUPPLEMENTARY TABLES

**Supplmentary Table S1**. The 17 columns in the final table.

**Supplmentary Table S2**. The portion of detectable bases in exonic regions by gene.

**Supplmentary Table S3**. The number of validated substitutions by KmerVC in TCGA samples for 1) Mutect2, 2) Varscan, 3) Muse, and 4) somaticsniper.

**Supplmentary Table S4**. The portion of Mutect2 subsitutions validated by KmerVC and portion of Mutect2 mutations identified by Varscan, Muse, and somaticsniper.

**Supplmentary Table S5**. The number of validated insertion/deletions by KmerVC in TCGA samples for Mutect2.

**Supplmentary Table S6**. The number of validated insertion/deletions at MS DNA by KmerVC in TCGA samples.
